# Hybrid quantum-classical de novo design of MHC-binding peptides

**DOI:** 10.64898/2026.07.09.736951

**Authors:** Emilie Sofie Engdal, Jonathan Funk, Omar Bacarreza, Laura Machado, Kristoffer Haurum Johansen, Janine Kemming, Thorin Farnsworth, Valentas Brasas, Rodrigue Yves Louis Lefèvre-Morand, Mateusz Slysz, Oliver Luscombe Nørregaard, Oliver August Dall’ Alba Sandberg, Alexander Makarovskiy, Peter Lodahl, Carlos G. Acevedo-Rocha, Krzysztof Kurowski, Sine Reker Hadrup, William R. Clements, Timothy P. Jenkins

## Abstract

Deep generative models have become a leading approach for designing therapeutic molecules, yet efficiently exploring vast biomolecular sequence spaces remains difficult, particularly for targets with limited training data. The prior distribution that seeds a generative model shapes which regions of sequence space it explores, and recent work suggests that non-classical distributions sampled from quantum processors can serve as a structured alternative to the factorised Gaussian priors used by default. Whether such priors help on complex biological design tasks has been largely untested. Here we present what is, to our knowledge, the first end-to-end hybrid quantum-classical pipeline for de novo design of MHC class I-binding peptides, coupling a generative adversarial network (GAN) to latent vectors sampled from a real photonic quantum processor. Tested in silico across 131 HLA alleles, quantum-derived priors increased the yield of predicted strong binders, with the largest relative gains for understudied alleles where classical baselines perform worst. We selected three understudied alleles for further evaluation, finding that large gains coincided with broader sequence exploration at non-anchor positions while anchor specificity was preserved. On these three alleles, we validated the designs in vitro using peptide-MHC stability ELISAs, confirming that quantum-designed peptides are potent stabilisers of peptide-MHC class I complexes. These results establish structured, hardware-realisable non-classical priors as a useful inductive bias for generative peptide design, with direct relevance to personalised immunotherapies and vaccines.

## 1 Main

Peptide-based vaccines and T-cell therapies depend on identifying short peptides that can be effectively presented by major histocompatibility complex (MHC) class I molecules to initiate protective immune responses. Only a small fraction of possible sequences form stable peptide–MHC (pMHC) ligands, and large-scale immunopeptidomics has shown that the mapping from amino acid sequence to antigen processing and presentation is structured, allele-specific and highly nonlinear [1–3]. Consequently, though state-of-the-art models achieve strong performance in predicting pMHC binding for many human leukocyte antigen (HLA) alleles [4, 5], the *de novo* generation of diverse, allele-specific peptide candidates remains challenging, particularly within sparse data regimes [6]. This limitation becomes especially apparent for understudied or structurally distinct alleles in which the empirical training distribution is biologically constrained. For example, HLA-A*31:01 is relatively rare in European populations but is strongly associated with severe T-cell mediated adverse drug reactions, underscoring the need to precisely delineate its peptide presentation landscape [7]. Generating novel binders for such alleles therefore provides a useful test case for novel methods that aim to navigate these complex sequence spaces where training data is biologically bottlenecked.

Generative models offer a promising route for exploring the vast sequence space underlying peptide design. Deep generative approaches can learn statistical structure from natural sequences and propose new candidates with desired biochemical properties [8–10]. In models that rely on mapping a “prior” or “latent” distribution to a data distribution, such as generative adversarial networks (GANs) and flow matching models, the prior is commonly chosen to be a simple classical distribution, typically isotropic Gaussian [11–13]. Emerging work suggests that the structure of the prior distribution can influence which regions of design space a model can access for biological applications [14], and that alternative priors may enable different or broader modes of exploration in high-dimensional landscapes [15, 16].

Quantum devices offer one such alternative. Photonic quantum processors effectively sample from high-dimensional, strongly correlated probability distributions that are not efficiently reproducible by classical algorithms [17–19]. Recent studies suggest that quantum-derived prior distributions can enhance the generative process in a range of tasks [20–22]. Bacarreza et al. (2026) [23] benchmarked GANs on small-molecule generation and found that photonic quantum priors yielded greater chemical diversity than alternative classical distributions. Ghazi Vakili et al. (2025) [24] hybridised a 16-qubit circuit with a classical LSTM to generate candidate inhibitors for the KRAS-G12D cancer target; their pipeline produced 15 compounds for synthesis, two of which showed inhibitory activity against KRAS in vitro. These examples motivate the hypothesis that quantum prior distributions can act as a useful inductive bias, increasing diversity in complex design tasks. To date, however, such demonstrations have been confined to small-molecule and image generation; whether non-classical priors transfer to biological sequence design, where the mapping from sequence to function is sparse remains untested.

We therefore reasoned that incorporating a quantum-derived prior distribution into a peptide generative model could systematically influence how allele-specific sequence spaces are explored, independent of model architecture or training objective. To isolate the effect of the prior, we adopted a conditional GAN framework which, while not the current state of the art in peptide generation, provides a controlled setting for modifying the prior distribution without confounding changes in model depth, conditioning strategy, or optimisation scheme [25, 26]. Our aim was to test whether structured, non-classical prior distributions obtained from quantum processors could shape peptide generation in ways relevant to MHC class I ligand discovery. To this end we built an end-to-end pipeline: we condition a GAN on latent vectors sampled from a real photonic quantum processor, generate peptide candidates across 131 HLA alleles, and validate selected designs in vitro by peptide-MHC stability ELISA for three understudied alleles.

## 2 A hybrid quantum-classical GAN model for MHC-binding peptides

To evaluate how quantum-derived priors influence peptide generation, we first required a large and well-curated set of experimentally validated pMHC ligands. We therefore used the curated ligand dataset assembled for NetMHCpan-4.1, which aggregates mass spectrometry-validated, naturally presented HLA class I ligands from the Immune Epitope Database [4, 27]. We restricted the dataset to 9-mers, the dominant length class of MHC class I ligands, to evaluate the effect of the prior distribution without confounding the analysis with variable-length modelling. This yielded 105,970 ligands (approximately 77,000 unique sequences) across 126 distinct MHC class I molecules (Figure 1a). This dataset provided both the training distribution and the conditioning labels for our generative model.

**Figure 1:**
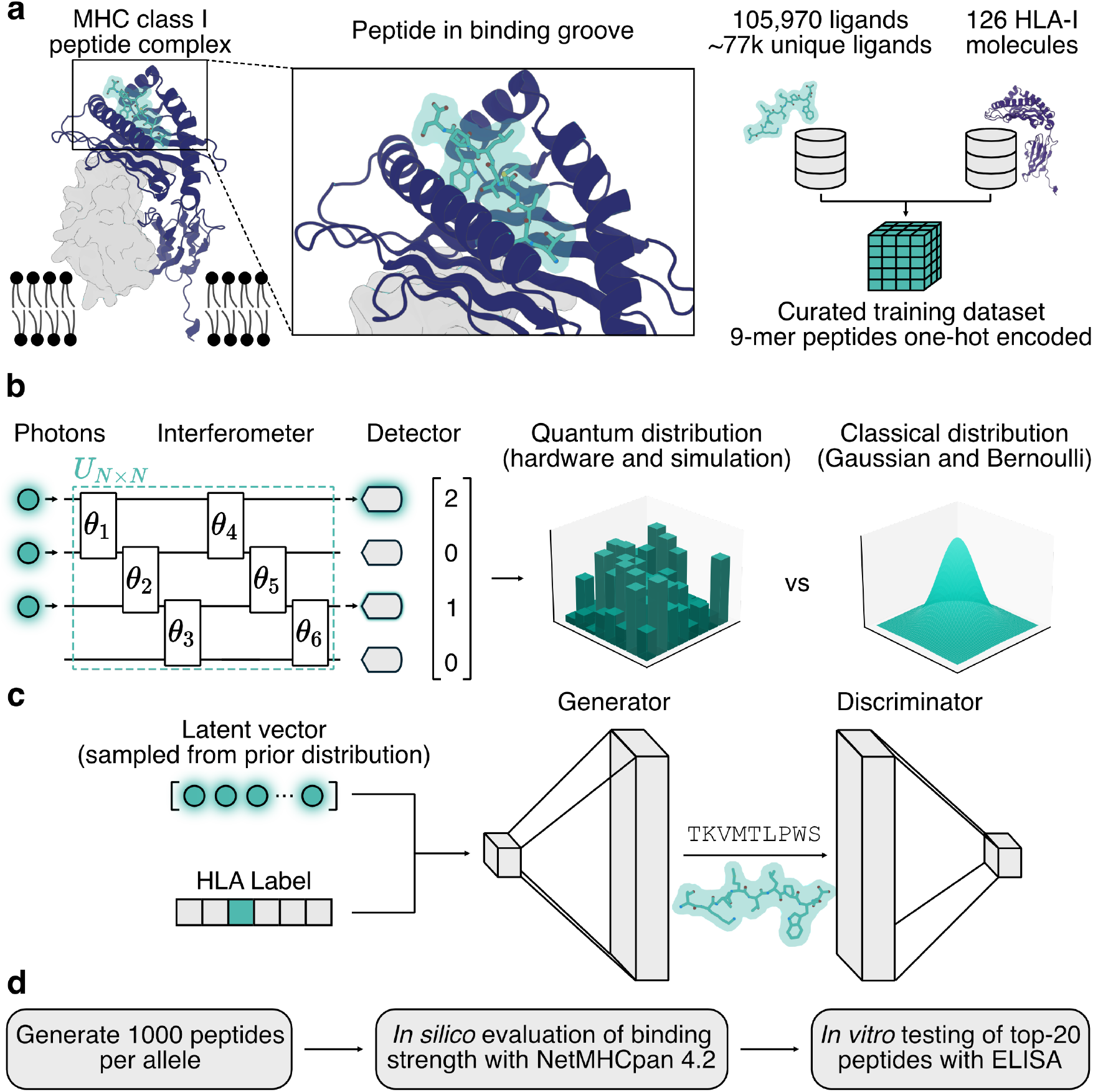
Overview of our workflow for hybrid quantum-classical de novo design of MHC-binding peptides.**(a)** Structural overview of a peptide bound within the MHC class I peptide-binding groove. We construct a dataset of 105,970 binding peptide-HLA pairs from the Immune Epitope Database, involving approximately 77,000 unique length-9 peptide sequences and 126 different HLA alleles. **(b)** We prepare a prior distribution using a 32-mode photonic quantum processor that implements a boson sampling protocol. As illustrated here with 4 modes, it operates by propagating multiple photons through a linear optical circuit composed of beam splitters, creating a superposition state. Photon number measurements at the output of each mode indicate where the photons left the circuit. Each measurement result can be interpreted as a sample from a probability distribution that is hard to simulate classically, in contrast to simple classical Gaussian or Bernoulli distributions that we use as baselines. **(c)** We use this distribution as a prior distribution in a generative adversarial network (GAN). In our GAN architecture, the generator is trained to map both latent vectors drawn from the prior distribution and an HLA label to peptide sequences. The discriminator is trained to distinguish between generated peptides and real peptides from the original dataset. **(d)** After training, we evaluate the trained generator by producing 1000 peptides per target allele. We then perform in silico evaluation of peptide binding strength using the NetMHCpan model. For three understudied alleles where large gains are observed, we evaluate the best 20 peptides in vitro using peptide-MHC stability ELISAs to verify their binding strength against the target allele.

We formulated the peptide generation task using a conditional GAN [11], chosen for its flexibility in allowing direct manipulation of the prior distribution without altering model architecture or conditioning. In this framework, a generator maps vectors drawn from a prior distribution to synthetic peptides, while a discriminator attempts to distinguish real ligands from generated ones. By conditioning both networks on HLA alleles, the GAN learns allele-specific sequence constraints present in the immunopeptidome.

Because our goal was to isolate the effect of the prior, we intentionally kept the GAN architecture simple and consistent across all experiments (Supplementary Figure 5 and Figure 6). The prior distribution is where we introduced variation. Conventional GANs typically draw latent vectors from factorised Gaussian noise because it is easy to sample from and empirically often achieves satisfactory results. However, theoretical and empirical work suggests that the structure of the latent prior can influence which solutions the generator can reach, particularly for multimodal targets or datasets with rich higher-order correlations [15, 28]. pMHC ligand sets are a natural example: anchor positions are tightly constrained while non-anchor positions vary widely, presenting exactly the kind of structured, correlated target where the choice of prior might matter. Although trainable or flow-based priors can approximate more complex distributions [12, 29], they introduce additional complexity that can confound attempts to attribute behavioural differences to the prior alone.

To test whether non-classical priors influence peptide generation, we replaced the Gaussian prior distribution typically used in GANs with distributions obtained from quantum processors. In general, the output distributions from quantum processors are highly correlated and cannot be classically factorised into independent components [30]. They therefore represent a qualitatively different prior from the uncorrelated Gaussian noise used in standard generative models. Motivated by recent theoretical and empirical studies suggesting that structured non-classical priors can improve mode coverage and alter generative exploration [20, 22, 23, 31–33], we sought to examine their impact in the context of peptide–MHC ligand design.

We generated latent vectors using both a real 32-mode photonic quantum processor and classical simulations of the same architecture. For the quantum processor, we used Gaussian Boson Sampling (GBS), a non-universal model for quantum computing [34]. In GBS, non-classical squeezed states of light are injected into a linear-optical circuit in which photons interfere with each other, creating an entangled quantum state. Photon-number-resolving detection then projects the state onto a discrete photon-number pattern, which serves as the latent vector (Figure 1b). In contrast to commonly used prior distributions, the output photon number patterns are in general correlated and non-uniform [35]. Photonic systems are well-suited for generating latent distributions because they avoid the decoherence-driven noise that tends to push output distributions in other quantum modalities close to uniform bit strings [22, 36]. We used a boson sampling system hosted at the UK National Quantum Computing Centre that uses a time-bin interferometer architecture [37, 38] and photon number resolving superconducting nanowire detectors. In addition to the hardware-generated samples, we produced simulated boson sampling distributions [39] to provide matched quantum-native priors while controlling for device noise.

Our experiments compare four prior distributions of identical latent dimension 32: two classical baselines (factorised Gaussian and Bernoulli), and two bosonic quantum distributions (a simulated Gaussian boson sampler and a real distribution measured on the photonic processor). In the Supplementary we also report results for a simulated bosonic non-quantum distribution with distinguishable photons, which do not interfere with each other. Using both real and simulated distributions in the subsequent experiments allowed us to assess the robustness of the approach to hardware imperfections as well as the contribution from the quantum distributions.

## 3 Performance of GANs trained with classical and quantum priors

We trained multiple conditional GANs on the curated pMHC ligand dataset, varying only the choice of prior distribution, and evaluated how prior choice affected binder yield across alleles of differing training-set frequency (Figure 2). To assess functional output, we generated 1000 peptide sequences across 131 HLA alleles (including 5 that were not in the training dataset) and predicted binding affinities using NetMHCpan

**Figure 2:**
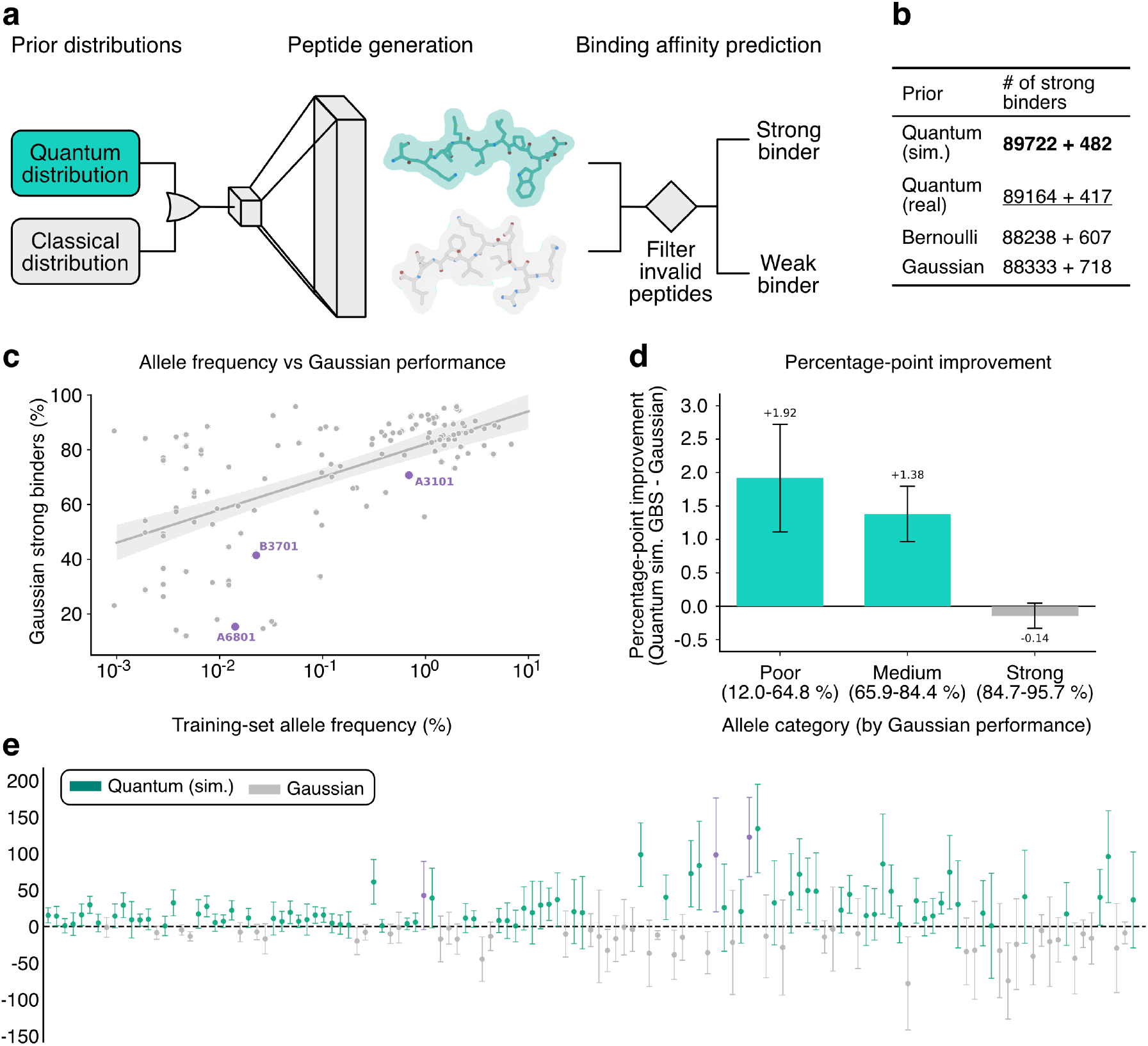
Evaluation of peptide-generating GANs using quantum and classical prior distributions. **a)** Evaluation pipeline. Generated peptide sequences were post-processed to remove invalid outputs, and binding affinity was assessed using NetMHCpan 4.2 rank score [5] to classify candidates as strong or weak binders. **b)** Number of strong binders produced by each prior, out of 1000 generation attempts for each of 131 target alleles. The quantum priors are found to produce improvements over the baseline classical Gaussian and Bernoulli priors. **c)** Percentage of strong binders produced by the Gaussian prior for each target allele, as a function of the frequency at which the allele occurs in the training dataset. Despite strong performance for the most common alleles, the model often struggles to produce strong binders for the least common alleles. **d)** Performance difference between the simulated quantum and Gaussian priors across different categories of alleles where the Gaussian prior achieves poor, medium and strong performance in terms of number of strong binders. The simulated quantum prior improves performance specifically on those alleles where Gaussian priors exhibit poor or medium performance. **e)** The difference in the number of predicted strong binders produced by simulated quantum versus Gaussian priors for each allele. Alleles are ordered by their frequency in the training data, with the rarest alleles on the right. The quantum prior outperforms the Gaussian prior on 63% of alleles, with large differences visible for several alleles. For **c** and **e**, the purple alleles are selected for further in-vitro study. For **b-e**, all results are averages over 30 training seeds, and error bars correspond to *±* one standard error of the mean.

4.2 [5]. Specifically, the eluted ligand (EL) likelihood scores were converted into percentile rank scores to enable cross-allele comparisons, accounting for inherent differences in binding affinity distributions between MHC molecules. Strong binders (SB) were defined using a percentile rank threshold of 0.5%. Each configuration was trained using 30 random seeds to reduce stochastic variability. To remove trivial distribution mismatches as a potential confounding factor, we rescaled the quantum-derived latent distributions to have zero mean and unit variance in each dimension.

First, we observe that the models conditioned on the quantum priors produced a larger number of strong binders compared to the Gaussian and Bernoulli priors (Figure 2b). The small difference (within one standard error) between the quantum priors from the simulation and the real system suggests that imperfections in the real hardware such as photon loss have at most limited impact on model performance. The Bernoulli and Gaussian priors performed similarly to one another and below both quantum priors, consistent with simple factorisable priors being less well matched to this task. Additionally, we found that the quantum distributions led to more consistent performance, with smaller standard deviations in the number of strong binders (see Supplementary).

While the average gain across all alleles is modest, it was concentrated in the alleles where the classical baselines performed worst and arguably an improvement matters the most. The number of strong binders produced varied widely across alleles (Figure 2c). While strong binders were consistently produced for the most common alleles in the training set, fewer strong binders were produced for more rare alleles. Comparison of the simulated quantum prior to the Gaussian prior (Figure 2d), suggested that the performance improvement concentrated among rarer alleles where the Gaussian prior exhibited poor (*<* 65% strong binders) or medium performance (65 − 85% strong binders). A head to head comparison between these two priors on the number of strong binders produced across all alleles is shown in figure 2e. The mean number of strong binders generated by the quantum prior was higher than Gaussian on 63% of alleles. While differences were modest for the more frequent alleles, they increased for the least frequent ones. A per-allele linear mixed model confirmed a statistically significant improvement from the quantum prior over the Gaussian baseline (simulated quantum: +10.6 strong binders per 1000, p = 0.001; real quantum: +6.3, p = 0.049; Supplementary).

Taken together, these results suggest that the quantum priors improved generalisation to data-poor regions of the sequence space. The ability to generalise in low data regimes can be of particular relevance in therapeutic contexts where rare or patient-specific alleles are often encountered. Experiments involving additional prior distributions, as well as further statistical analysis, can be found in the Supplementary.

## 4 Allele-specific dynamics in data-sparse regimes

To understand how the priors influence allele-specific behaviour, and to guide peptide selection for down-stream in vitro testing, we selected three HLA alleles (Figure 2). HLA-A*68:01, HLA-B*37:01 and HLA-A*31:01 all appeared with low frequency in the original training set, where the Gaussian baseline under-performed compared to the general trend line (Figure 2c), and where the quantum priors outperformed the Gaussian prior. The chosen alleles occur at low-to-moderate frequency in European populations (HLA-A\**68:01, 3*.*6%; HLA-B\**37:01, 1.4%; HLA-A*31:01, 2.3%; allele frequencies from a German donor cohort of 3.46 million individuals [40]). HLA-A*68:01 belongs to the A3 supertype and shares its canonical anchor grammar, a small or aliphatic residue at P2 and a basic residue (Arg/Lys) at the C-terminus, even though it is itself underrepresented in European immunopeptidomics datasets relative to common A3 members such as HLA-A*03:01 and HLA-A*11:01 [40]. Because this anchor motif is well represented across the supertype, sequence-based predictors generalise to it reliably. HLA-B*37:01 is the opposite case: it carries an unusual acidic P2 anchor (Asp/Glu) shared by only a small subset of HLA-B alleles [41]. This feature is sparsely represented in training data and sits atypically among the predominantly hydrophobic-anchor HLA-B repertoire. These alleles therefore bracket the regime we care about, a well-supported target where prediction is easy and a sparsely-supported one where it is not.

We compared the training curves for these alleles and studied the diversity of the generated peptides (Figure 3). The training curves for HLA-A*31:01 and HLA-A*68:01 showed a consistent gain throughout training for both real and simulated quantum priors. For HLA-A*68:01 the quantum priors produced roughly twice as many strong binders as the Gaussian prior across training. The persistence of this advantage throughout training for both alleles is consistent with the interpretation that the quantum priors endowed the model with a useful bias for producing strong binders for those alleles. The training curves for HLA-B*37:01 are noisier. Though the results for HLA-B*37:01 suggest an advantage from using quantum priors, stochastic variation between seeds cannot be ruled out.

**Figure 3:**
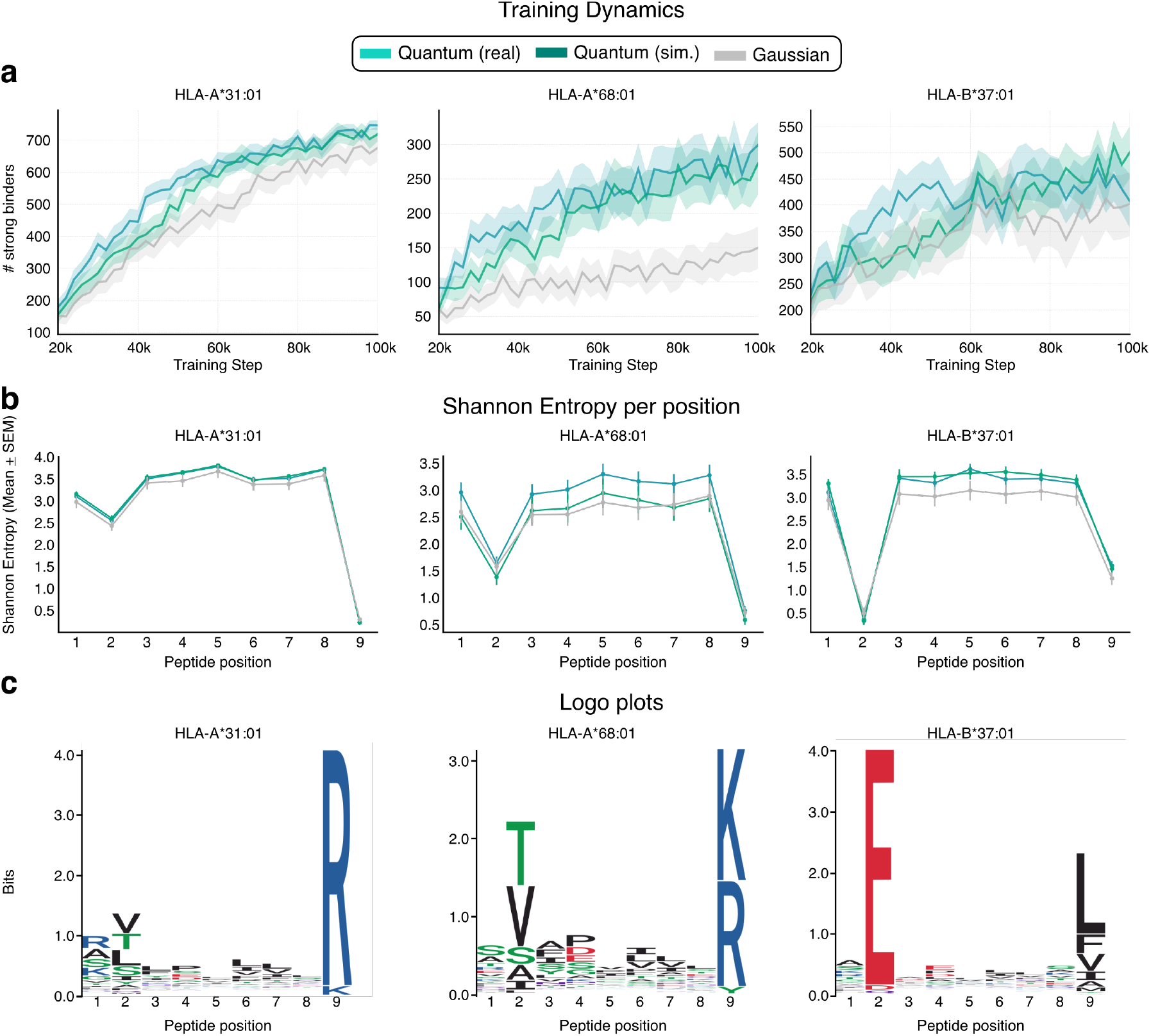
Allele-specific effects of quantum priors on binder yield and sequence diversity. **a)** Mean number of strong binders produced out of 1000 generation attempts at different steps during training, for different priors and the three selected HLA alleles. **b)** Positional diversity of generated binders. Mean Shannon entropy across peptide positions for the three HLA alleles, comparing prior types. Higher entropy indicates greater amino-acid diversity at a given position, reflecting broader exploration of allele-compatible sequence space while maintaining predicted binding. In **a** and **b**, error bars correspond to *±* one standard error of the mean across 30 training seeds. **c)** Sequence logos of strong-binding 9mer peptides generated from Quantum (real) latent distribution for alleles A*31:01, A*68:01 and B*37:01.

To assess whether the binder-yield differences reflect broader exploration of allele-compatible sequence space rather than repeated sampling of the same motifs, we quantified positional sequence diversity by computing the Shannon entropy of amino acid usage at each peptide position across the generated sequences (Figure 3b). Models using quantum priors produced peptides with higher overall positional entropy than the Gaussian prior, and this increase concentrated at non-anchor positions while anchor positions remained constrained. The generated motifs recovered the known anchor grammar of each allele, the basic C-terminus of HLA-A*68:01 and the acidic P2 of HLA-B*37:01 (sequence logos, Fig. 3), confirming that the added diversity reflects broader exploration of permissible sequence space rather than degraded anchor specificity. A well trained model should retain low entropy at P2 and P9, where allele-specific anchor preferences constrain binding, and gain entropy primarily at non-anchor positions where sequence variation is biologically permissible. Elevated entropy at anchor positions would instead indicate less specific rather than more diverse generation. For HLA-B*37:01, the consistently higher diversity achieved in the non-anchor positions using the quantum priors indicates underlying differences between the quantum and Gaussian priors that are not captured by the training curves alone. Across all three HLAs, the greater diversity provides a plausible explanation for the improved binding predictions, as a wider non-anchor search space increases the likelihood of identifying high-affinity motifs.

## 5 In vitro validation

Since classical predictors are known to lose precision on rare alleles [6], predicted binding alone is insufficient and direct experimental validation is required. We tested designs for the three alleles identified above: HLA-A*68:01 and HLA-A*31:01, which carry well-supported anchor motifs, and HLA-B*37:01, whose atypical acidic P2 anchor is sparsely represented in the training data. Using the configuration in which the prior was sampled from the physical photonic quantum processor, we selected the 20 highest-ranked peptides per allele by EL %rank for synthesis. As negative controls we selected, per allele, the three peptides with the lowest predicted binding across all priors. All tested sequences and predicted scores are listed in Supplementary Table 8. Following UV-mediated peptide exchange, pMHC integrity was quantified by sandwich ELISA as blank-subtracted absorbance at 405 nm (OD405), with positivity defined relative to the UV-only no-peptide control (mean + 3 s.d.; dashed line in Figure 4b).

**Figure 4:**
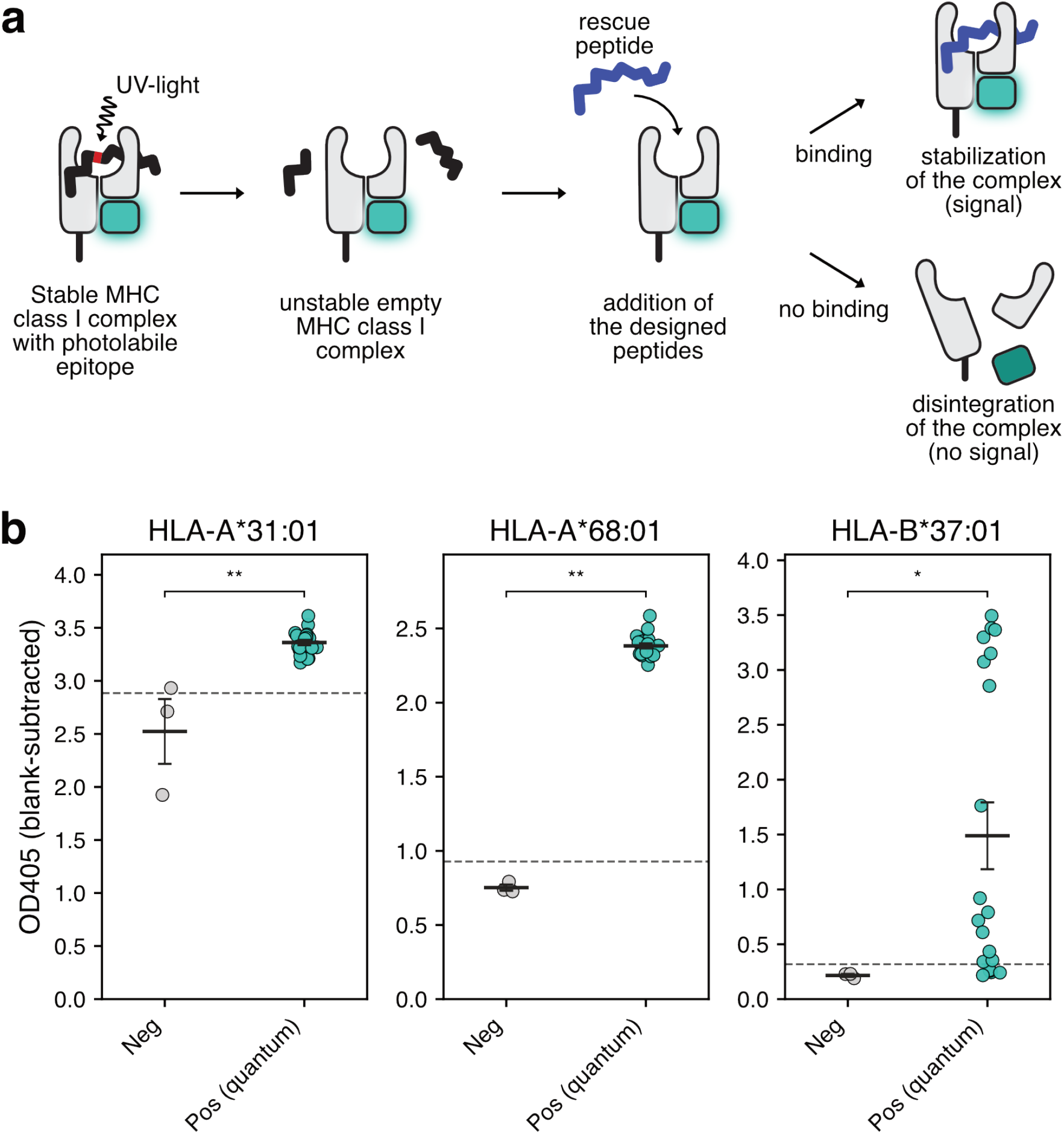
In vitro stabilisation of pMHC complexes by peptides generated with a quantum-derived prior. **a)** Schematic of UV-mediated ligand exchange. UV irradiation cleaves a photolabile conditional peptide within a pre-formed MHC class I complex, yielding a peptide-receptive intermediate that rapidly destabilises unless a “rescue” peptide is supplied, in which case stable pMHC formation is recovered and quantified by ELISA (see Methods; adapted from established UV-exchange protocols). **b)** Peptide-dependent pMHC stability measured by sandwich ELISA after UV exchange for the three selected alleles. Points represent the mean of technical triplicates; horizontal bars indicate the mean *±* s.d. The dashed line indicates the positivity threshold defined as the mean + 3 s.d. of the UV-only control without peptide.

**Figure 5:**
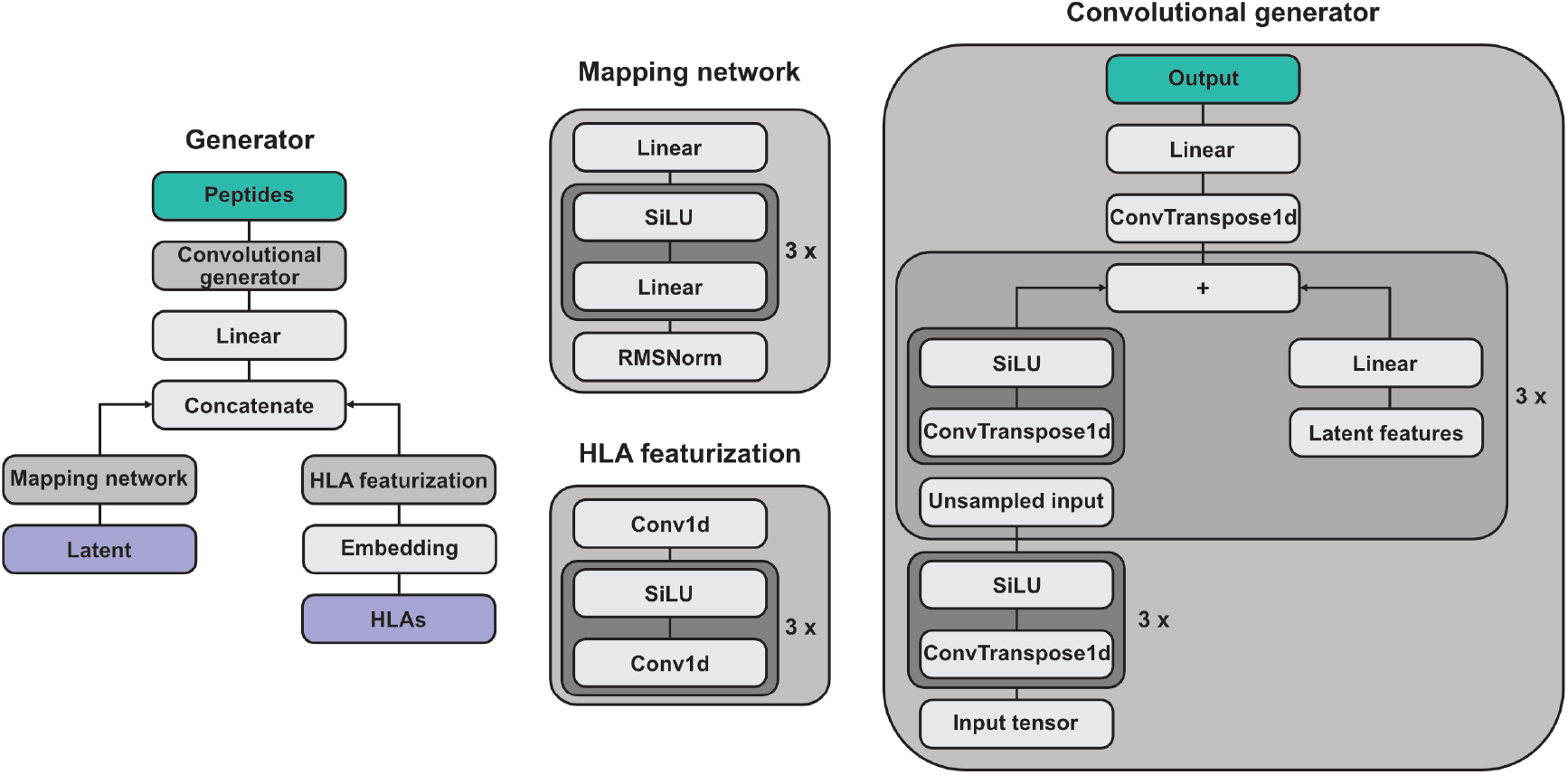
Generator architecture.

**Figure 6:**
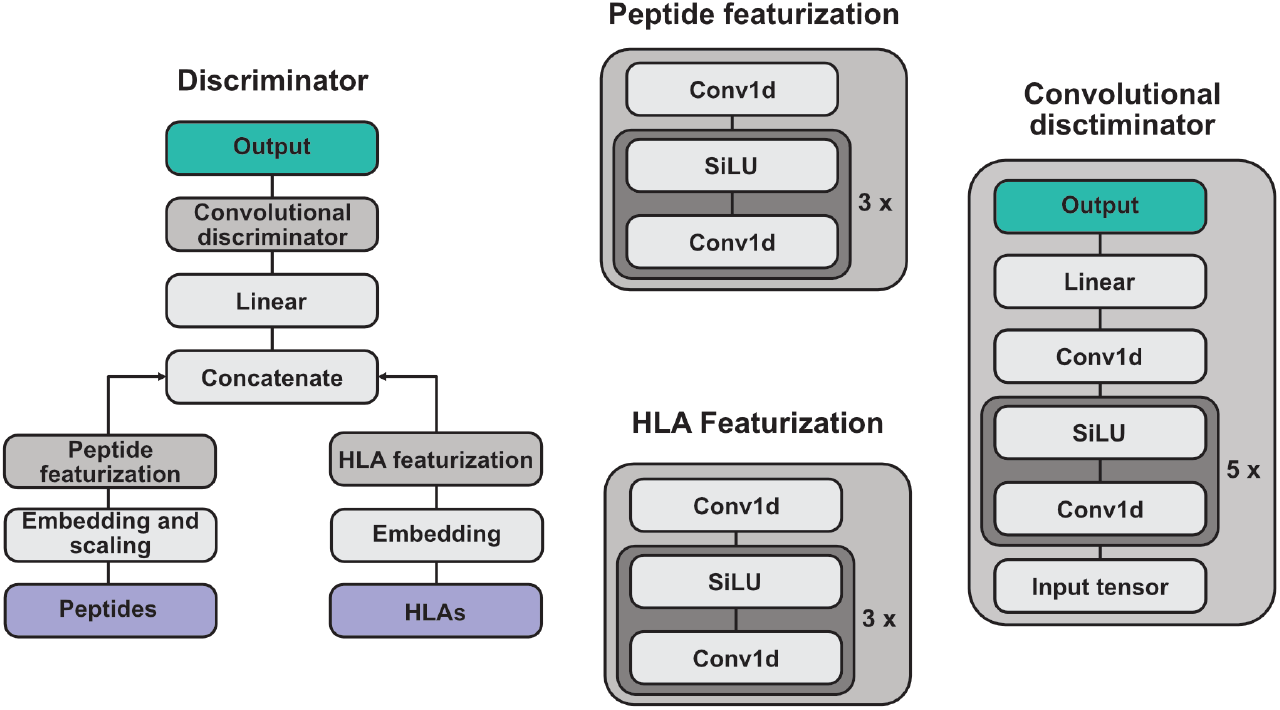
Discriminator architecture.

Across all three alleles, quantum-prior-generated peptides produced OD405 signals well above both the negative controls and the positivity threshold, confirming that they stabilise peptide-MHC class I complexes in vitro (Figure 4b). All three negative-control peptides per allele failed to stabilise the complex. For HLA-A*31:01 and HLA-A*68:01 all tested predicted binder peptides exceeded the positivity threshold. The dynamic range across positives was particularly narrow for HLA-A*68:01 (OD405 ∼ 2.25-2.6) indicating a uniformly successful validation. For HLA-B*37:01 the quantum-prior peptides include many strong binders but also several failures. This result underscores the need for in-vitro validation for rare alleles where classical predictors can be less reliable. The wide range of binding strengths is consistent with the high sequence diversity observed for HLA-B*37:01 which has less well-defined anchors than HLA-A*68:01, and suggests that a broader sequence exploration space may capture high-affinity outliers at the cost of greater variance in individual peptide performance. We note that empty/open HLA-B*37:01 is conformationally sensitive, with chaperone binding inducing detectable conformational changes in the B*37:01/*β*2m complex [42], which may further contribute to the wider spread of in vitro outcomes for this allele. In the Supplementary, we additionally compared the best quantum-generated peptides against the best classical-generated peptides for these alleles. Though the mean OD405 values were similar between sets, the strongest single binders for each allele was produced by the quantum-prior model. Taken together, these results demonstrate that peptides derived from a quantum prior using a real quantum processor can successfully bind to MHC-I complexes across understudied alleles.

## 6 Discussion

In this work, we investigated how the choice of prior distribution influences generative modelling of MHC class I peptide ligands. By replacing the commonly used factorised Gaussian prior with structured, quantum-derived prior distributions, we observed consistent differences in training dynamics, sequence diversity and binder yield, with the largest gains concentrated in alleles where the classical baseline underperforms. In vitro validation across three alleles confirmed that quantum-prior peptides reliably stabilise pMHC complexes, including for HLA-B*37:01, the allele on which sequence-based predictors are least reliable. The greater variance observed for HLA-B*37:01, in both training dynamics and in vitro outcomes, is most parsimoniously attributed to its atypical acidic P2 anchor being sparsely represented in the training data, which reduces predictor reliability across all priors, rather than to a property of the quantum prior. Notably, the generator still recovered the correct acidic-anchor motif and produced experimentally validated binders even for this hardest allele. Together, these findings suggest that the prior distribution acts as a meaningful inductive bias in peptide generative models, shaping how allele-specific sequence space is explored rather than redefining the underlying limits of MHC binding prediction.

A natural question is whether similar gains could have been achieved without a quantum processor. While the scale of probability distribution considered here is classically simulable, at larger sizes an exact simulation is intractable. These larger sizes may be required for larger-scale generative models, where prior distribution sizes larger than 32 are often required. At those larger scales, approximate simulations methods still exist [43–45], but they may not be suitable for this algorithm. These approximate methods correctly reproduce the first few moments of the distribution and accurately describe the statistics of a realistic lossy photonic systems. However, the wall-clock times for all these simulation methods are significantly longer than those of a physical system, which is likely to make them impractical for the algorithm considered here. An analysis of runtimes can be found in the supplementary materials. A related question is whether other types of classical distribution beyond the ones considered here could have produced a similar improvement. While such classical distributions likely do exist, the problem of finding the optimal latent distribution is in general hard [15]. Selecting a quantum distribution for its highly correlated and non-factorisable nature, which we hypothesised could be beneficial, provides justification for selecting a quantum prior distribution a priori. We emphasise that these results do not constitute a demonstration of quantum advantage: the system sizes used here remain classically simulable, and the measured gains over classical priors, while consistent, are modest. Our claim is the narrower one that a structured, non-classical prior sampled from real quantum hardware functions as a useful and hardware-realisable inductive bias for generative peptide design.

Extensions of this work could include adapting this approach to state-of-the-art architectures such as flow matching models [46, 47], which similarly rely on prior distributions and may benefit from non-classical priors [23]. Second, though peptide–MHC binding is a prerequisite for T cell recognition, it is not equivalent to immunogenicity. True immune activation depends on additional factors such as peptide processing, TCR availability, immune context and conformational dynamics [48]. Future generative models could thus benefit from extending their objectives beyond predicted MHC binding to include features that approximate molecular dynamics and structural adaptability. Quantum prior distributions may offer an opportunity in this context, helping design peptides that are not only strong MHC binders but can also elicit effective immune responses.

The pronounced effects observed in data-sparse alleles also raise the possibility that allele-conditioned generative models could help address uneven training-data coverage across HLA alleles. Because modern pan-allelic predictors improve as additional allele-resolved binding and ligand data become available [5, 6], a practical future direction would be to test a retraining loop in which experimentally validated, model-generated binders for low-frequency alleles are added to the training set, followed by evaluation of whether cross-validation performance improves specifically in sparse-allele regimes.

Our work indicates that the structure of the prior distribution can influence how efficiently generative models capture the allele-specific constraints underlying MHC class I peptide binding. By integrating a photonic quantum processor as a source of structured prior distributions within an otherwise classical generative framework, and validating the resulting peptides experimentally, we demonstrate how quantum-derived priors can act as a useful inductive bias for modelling peptide–MHC binding landscapes. Such approaches may contribute to the efficient exploration of binding-compatible sequence space, particularly for alleles that are poorly served by current training data. As quantum hardware and hybrid modelling approaches mature, these techniques could be incorporated into broader immunoinformatics pipelines that account for antigen origin, expression, and downstream immunological constraints.

## 7 Acknowledgements

First we would like to thank Sofie Lindskov Hansen, who inspired this multi-institutional collaboration and enabled this project coming together. We would also like to thank David Jenkins for his feedback and helpful suggestions throughout this project. Finally, we would like to thank our funders. T.P.J, C.G.A.R and J.F. thank the Novo Nordisk Foundation Center for Biosustainability (NNF20CC0035580) who helped support this work. C.G.A.R. thanks “The Novo Nordisk Foundation” NNF BRIGHT Grant number: NNF24SA0100980. Further, this research has been supported by the Program of the Polish Ministry of Science and Higher Education “Applied Doctorate” realised in years 2022-2026 (agreement no. DWD/6/0142/2022), by the Poznan University of Technology (project no. 0311/SBAD/0746), and by ń Supercomputing and Networking Center Collaboration Agreement (project no. 323/PCSS/2024) with ORCA Computing Limited for Development, Testing and Benchmarking.

## 8 Author Information

### Authors and Affiliation

Correspondence and requests for materials should be addressed to W.R.C and T.P.J.

### Contributions

E.S.E. led the numerical experiments. J.F., O.B., L.M., T.F., V.B. and R.Y.L.L-M. all designed and implemented generative models and software used at different stages in this project. K.H.J., S.R.H., and J.K. designed and implemented the in-vitro experiments. M.S. implemented experiments involving quantum photonic hardware. O.L.N., O.A.D.A.S and A.M. contributed to analysing the data and methodology. P.L., C.G.A.R., K.K. provided supervision for the project. W.R.C. and T.P.J. conceptualised and supervised the project. E.S.E drafted initial figures and manuscript. J.F. prepared the final figures, and all authors contributed to writing the manuscript.

Supplementary Information is available for this paper.

## 9 Code and Data Availability

Our code is available at https://github.com/orcacomputing/pep-q-gan. Note that the NetMHCpan executable and the samples from the quantum processor are proprietary components that are not included in the code.

## 1 Methods

### 1.1 Training set

The training data consists of 105,970 experimentally validated peptide ligands (∼77k of which are unique) of length 9 (9mers) binding to 126 distinct MHC molecules. The binders are obtained from mass spectrometry (MS) elution data made available through the Immune Epitope Database (IEDB) as described in NetMHCpan-4.1 [4, 27]. As opposed to conventional binding affinity data, MS peptidome data includes the context of antigen processing and presentation as it reflects peptides naturally presented on the cell surface [49].

### 1.2 Conditional hybrid quantum-classical WGAN-GP architecture

We use a conditional Wasserstein GAN with gradient penalty [50] (cWGAN-GP). Our cWGAN-GP architecture consists of three main components: the Generator (*G*), the Discriminator (*D*) and a prior distribution. We consider prior distributions drawn from either classical and quantum bosonic distributions as well as standard classical distributions (Gaussian and Bernoulli). All protein and peptide sequences are represented as one-hot encoded embeddings, where each bit represents an amino acid in a vector of 1s or 0s. Conditional frameworks for both *G* and *D* support targeted exploration of a biologically relevant sequence space, which are particularly important in molecular design tasks such as drug discovery or neoantigen generation.

#### 1.2.1 Generator and Discriminator

The Generator is a conditional generator that fuses information from a latent vector and a biological sequence (HLA), using embeddings, convolutional neural networks (CNNs), and fully connected layers, to produce novel peptide sequences. The Discriminator is a conditional classifier that uses deep convolutional layers to jointly process peptides and HLA pseudosequence information, outputting a score that indicates whether the input pair is real or generated. A full description of these models can be found in the supplementary materials.

### 1.3. Training

We used the same hyperparameters for all experiments: a batch size of 256, the AdamW [51] optimiser with a learning rate of 10^−4^, and 100000 training iterations. These hyperparameters were selected based on prior work in [23], and after finding that these led to successful model training and a high number of strongly-binding peptides we did not further optimise them.

### 1.4. Prior distributions

The latent vectors ***z***, which are all of size 32, act as an input to the Generator. Gaussian random noise and the Bernoulli distribution represent common classical baseline models for comparison. We also use three bosonic distributions: one simulated classical distribution from a Gaussian boson sampler with distinguishable photons, one simulated quantum distribution from a Gaussian boson sampler with indistinguishable photons, and one real boson sampling distribution measured from an ORCA Computing PT-2 system. Under mild complexity assumptions, the resources required to classically simulate a Gaussian boson sampler scale exponentially with the number of photons, making these simulations intractable beyond a few tens of photons [34, 39]. For each model, to sample from these distributions we first precomputed a dataset of samples from which we drew batches as required by the training process. Further information can be found in the Supplementary.

### 1.5. Model performance

At the end of each epoch, the model is evaluated by generating 1000 peptides for 131 alleles. Note that 5 of these alleles were not in the training set. Peptides with a length less than 8 were filtered out. To more accurately assess generative performance, the models were evaluated using NetMHCpan 4.2 predicted peptide–HLA binding rank score. Specifically, the eluted ligand (EL) likelihood scores were converted into percentile rank scores using NetMHCpan to enable cross-allele comparisons, accounting for inherent differences in binding affinity distributions between MHC molecules [5]. Strong binders are defined as percentile rank threshold of 0.5%.

### 1.6. Computational resources

The computational resources used for our experiments can be divided into the time taken to collect samples from the prior distribution and the training time.

It took 40 minutes to collect 500k samples with an ORCA PT-2. For the simulated Gaussian boson sampler size-32 models, 30 different sets of optical circuit parameters were used to precompute 1 million samples (a total of 30 million). The time taken to obtain 1 million samples using a simulated Gaussian boson sampler was 511. *±* 55 min 35.33 min. The time taken to collect samples from simulated distinguishable photons was negligible.

Each training experiment ran on a single NVIDIA HGX™ A100 GPU 80GB. The models were trained using the precomputed samples (for the quantum and distinguishable priors), and samples generated on the fly for the Gaussian and Bernoulli distributions. It took around 400 minutes to train each model.

### 1.7 Peptide–MHC class I UV exchange and stability ELISA

HLA-A*31:01, HLA-A*68:01, and HLA-B*37:01 monomers carrying a UV-cleavable conditional ligand were produced as described previously [52, 53]. UV-exchange and stability ELISA were performed with synthesised peptides (SB peptides) as previously described [52]. Briefly, MHC integrity following UV exchange was assessed by sandwich ELISA. ELISA plates were coated with 2 µg/ml Streptavidin (room temperature overnight), followed by attachment of UV-exchanged avitagged pMHCs, and staining with 1.4 µg/ml HRP-conjugated anti-*β*2M (clone 2M2, BioLegend). The HRP reaction was stopped using an ABTS stop solution after 10 minutes, and absorbance was measured at OD405 on an ELISA plate reader. Each peptide was assayed in n = 3 wells. A peptide was classified as positive when OD405 exceeded the UV-only no-peptide control threshold (mean +3SD). Peptide-dependent stability was quantified as the OD405 after subtracting the blank. Signals were evaluated relative to a UV control without peptide.

## A Supplementary materials

### A.1 Full model descriptions

#### A.1.1 Generator

##### Mapping Network

Inspired by StyleGAN [54], we use a mapping network that transforms the initial prior distribution using a small-scale trainable neural network. This mapping network consists of a multi-layer perceptron (MLP), and non-linear transformations (SiLU activations). This network creates intermediate “latent features”.

##### HLA Embedding and Featurisation

HLA indices are embedded and processed through a series of three 1D convolutional layers with SiLU activations and a final 1D convolution, hereby compressing all amino acid positions into a single feature vector capturing information from the entire sequence.

Latent features and HLA features are concatenated and projected, using a linear layer, to the size used by the convolutional generator.

##### Convolutional Generator

This is the main component of the generator. This module processes the combined features to generate the final output peptides. It uses transposed convolutions and latent injection at each stage to generate a complex, high-dimensional output from a lower-dimensional feature vector. Specifically, it is composed of a sequence of i) generating blocks: each block consists of transposed convolutional blocks and activations, and different numbers of input and output channels that progressively upsample and transform the input, ii) a transposed convolutional layer and iii) a final linear layer that outputs logits. The latent vectors are injected by mapping the latent features into shapes compatible with each generating block. At each stage (except for the first block), latent features are added to the previous outputs of the generating blocks. The logits are then passed through the softmax activation to provide discrete outputs (one-hot encoded peptide sequences) and differentiable sampling while maintaining gradients.

#### A1.2 Discriminator

##### Input Scaling and Peptide Featurisation

The peptide input is scaled by a learnable parameter, allowing the model to adaptively weight each input channel. After scaling, the peptide is processed through a series of two 1D convolutional layers with SiLU activations and a final convolutional layer, extracting hierarchical features capturing position specific dependencies (e.g. P2 and P9) in a motif-context.

##### HLA Embedding and Featurisation

HLA indices are embedded and passed through their own series of three 1D convolutional layers, also with SiLU activations and a final convolutional layer.

Peptide features and HLA features are concatenated and projected into a hidden size using a linear layer.

##### Convolutional Discriminator

The discriminator processes the combined features to produce a single discrete output score, indicating whether the input peptide is real or generated. It is composed of a stack of discriminating blocks, each consisting of a 1D convolution, and SiLU activation, followed by a convolutional layer that reduces the output to a single channel and a final linear layer that maps the output to a single scalar value.

#### A.1.3 Discriminator (critic) update

For each training iteration, the discriminator is updated 5 times before each generator update. Real peptide sequences ***x*** together with conditional HLAs **mhc** are drawn from the dataset, and generated peptide sequences 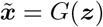 are produced by the generator and all are passed through *D*.

The discriminator computes scores for both real and generated peptides. The Wasserstein loss is calculated as the difference between the mean scores for real and fake samples. Then, gradient penalty is computed to enforce the Lipschitz constraint.

The total discriminator loss is the sum of the Wasserstein loss and the gradient penalty.

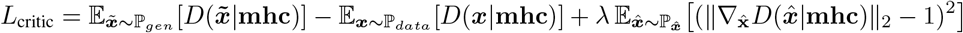

where 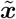 are the generated samples, 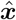 are random interpolations between real and fake samples, and *λ* is the penalty coefficient.

#### A.1.4 Generator update

After 5 discriminator updates, the generator is updated once. The generator produces peptide sequences 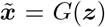 using the sampled latent vectors ***z*** ∼ ℙ_***z***_ and conditional HLAs **mhc** drawn from the dataset. The discriminator evaluates these generated samples. The generator loss is:

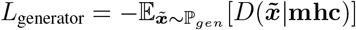

#### A.1.5 Sampling from quantum distributions

The two simulated distributions considered ideal lossless experiments and used single mode squeezed state inputs with an average of 0.5 photons per mode, such that the output measurements have an average of 16 photons in 32 modes. The simulation with indistinguishable photons was performed using the algorithm of [39], while the simulation with distinguishable photons reported in the supplementary materials was performed by summing the output statistics obtained by sequentially injecting each input state individually into the interferometer. To match the real quantum processor, the simulations considered time-bin interferometer circuits [38] consisting of a sequence of two identical delay lines. The beam splitters in these circuits were initialised randomly, and a dataset of 1 million samples per circuit was obtained for GAN training. We used different randomly selected beam splitter parameters for each seed.

The real quantum system that we used is the ORCA Computing PT-2 system deployed at the UK National Quantum Computing Centre, illustrated in Figure 7. Similar to the simulations, this system implements a time-bin interferometer architecture with two sequential delay lines. The delay lines are coupled to the “straight-through” channel via beam splitters with a splitting ratio that is a tuneable parameter. This parameter can be independently configured between time bins. Output photons are measured using photon number resolving superconducting nanowire detectors. Reaching a regime with an average of 16 photons in 32 modes is impractical due to optical losses and limitations to the squeezing parameter, so instead we used post-selection to only return results with at least 16 photons in 32 modes. This leads to an effective average photon number of 17.3. We randomly initialised the beam splitter parameters and collected a dataset of 500,000 samples to use for training. These same samples were used for all the 30 training seeds, which we note may lead to less variability between different seeds than the experiments with the simulated distributions.

**Figure 7:**
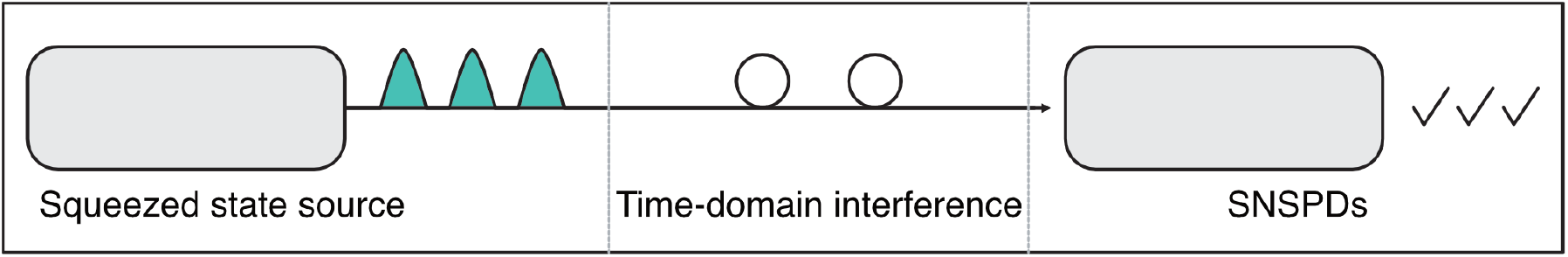
Quantum processor used to produce the quantum prior distribution. A quantum light source produces quantum states of light at regular time bins. These are sent into an interferometer consisting of a sequence of two identical delay lines, where the length of each delay line is such that each time bin interferes with subsequent time bins. This interference process produces an entangled quantum state. A photon number resolving superconducting nanowire detector (SNSPD) then performs a measurement collapsing the state and returning the number of photons measured in each time bin.

### A.2 Additional results with a distinguishable photon distribution

We also compared the quantum priors to another “distinguishable” classical prior consisting of a simulation of the photonic device in which all photons were no longer treated as identical. This distribution differs only from the quantum distributions in that distinguishability between photons removes the quantum interference that gives rise to non-classical correlations [55]. Our experiments yielded a small performance advantage in favour of the quantum prior, which is qualitatively in line with prior work [23]. Though these results are not conclusive, they are suggestive that this improved performance is connected to the specifically quantum properties of the prior.

### A.3 Additional analysis of the results

Here, we provide further analysis of the comparison between models trained with different prior distributions.

Figure 8 shows violin plots of the distribution of the number of strong binders produced by each approach, showing several qualitative differences. First, as reflected in table 1, the quantum distributions produce more strong binders on average. Second, they also have significantly lower standard deviation, showing that they produce strong binders more consistently.

**Table 1:**
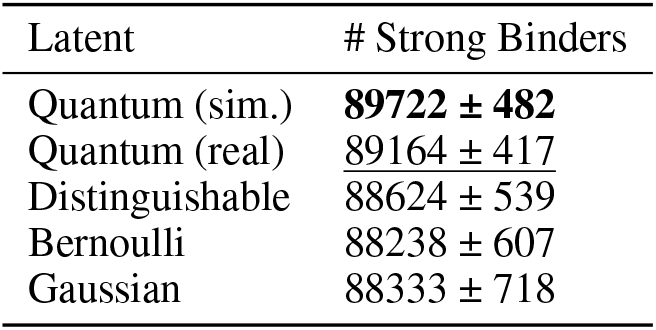
Summary Table (Mean ± SEM)

**Figure 8:**
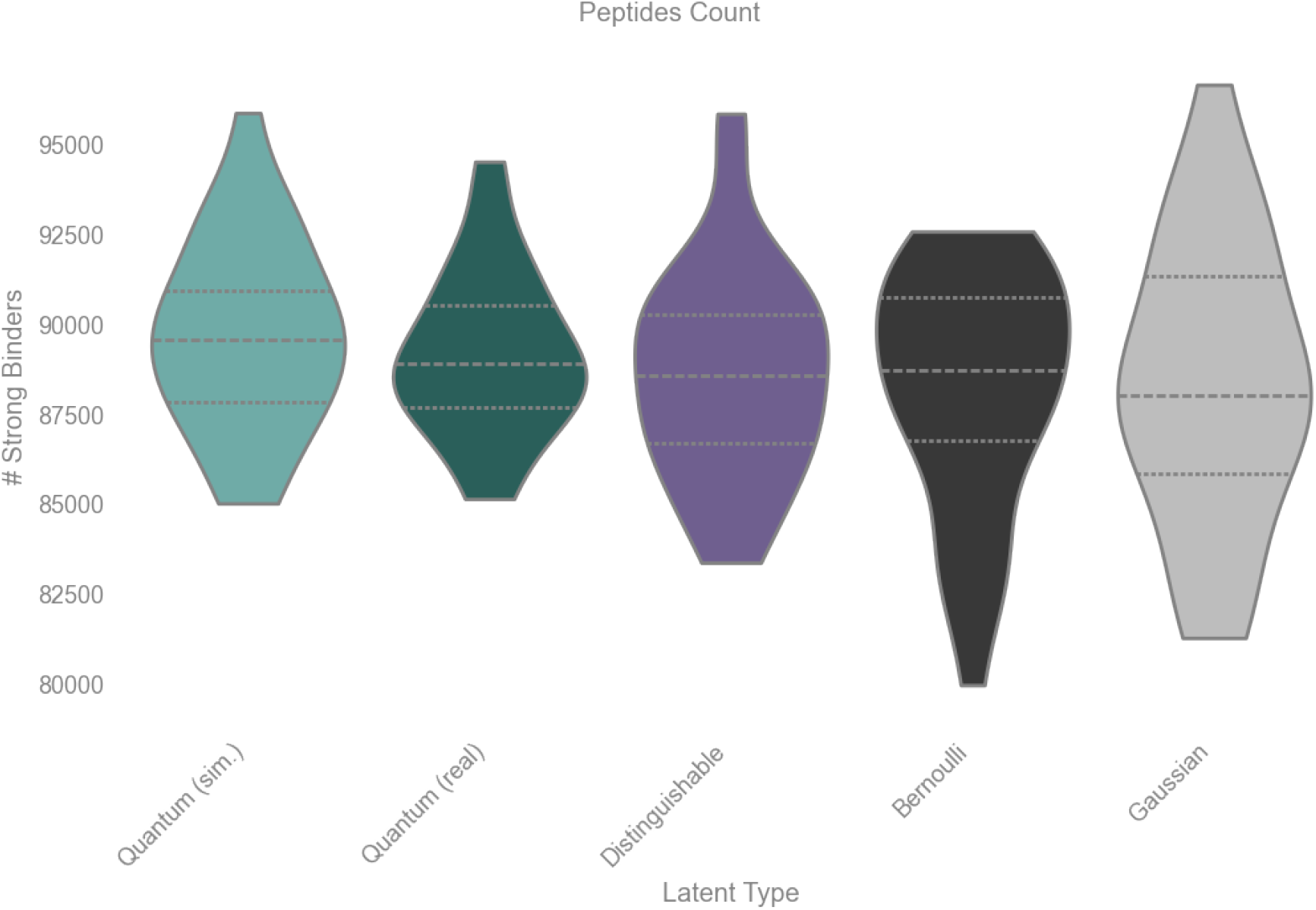
Distribution of strong binders for different prior distributions, estimated from 30 training seeds. The dashed line represents the median, and the dotted lines represent the quartiles.

The following sections discuss the statistical significance of the observed differences between the distributions. To report the significance of the difference in means, we use both the parametric T-statistic and the non-parametric Wilcoxon test. Given the visibly non-Gaussian nature of the distributions in figure 8, we expect that the Wilcoxon test is the most appropriate metric. Since our driving hypothesis, based on prior work such as [23], was that the quantum distributions would produce a larger number of strong binders, for the mean number of strong binders we report one-sided P values. To report the difference in standard deviations, for which we had no a priori expectation, we use the (two-sided) F-test and Levene test.

We also report statistics from a linear mixed model on the per-alleles differences between the priors. Compared to the mean and standard deviation of the aggregate number of strong binders which provide a coarse view of the differences in performance beyond the model, the mixed linear model returns the average expected improvement from using one prior instead of another on a randomly selected allele. Since this metric has not been used in prior work comparing quantum to classical priors, we report two-sided P values.

In the following, P values below 0.05 are reported in bold.

#### A.3.1 Comparison between Quantum and Bernoulli

Table 2 reports the statistics of the difference in mean and standard deviation of the number of strong binders produced by the simulated and real quantum distributions compared to the Bernoulli distribution. We find that the simulated quantum distribution significantly outperforms the Bernoulli distribution in the mean number of strong binders. The results for the real quantum distribution are only suggestive of a performance advantage in the mean number of strong binders. The results for the standard deviation for both distributions also suggest that the quantum distributions are narrower, in particular for the real quantum prior.

**Table 2:** Statistics - Quantum vs Bernoulli.

| Statistic | P-value (quantum sim vs Bernoulli) | P-value (quantum real vs Bernoulli) |
| --- | --- | --- |
| *Mean* |  |  |
| T-statistic | **3.04e-02** | 1.07e-01 |
| Wilcoxon | **4.81e-02** | 1.40e-01 |
| *Variance* |  |  |
| F-test | 1.11e-01 | **2.37e-02** |
| Levene | 2.43e-01 | 5.19e-02 |

Table 3 reports the results from the mixed linear model. On average, the Bernoulli prior produces 673.6 strong binders out of 1000 generation attempts for each allele. The simulated quantum prior produces 11.3 more strong binders than Bernoulli on average. The improved performance is statistically significant (z=3.5, p<0.001), with a 95% confidence interval of [5.0, 17.7]. The real quantum prior produces 7.1 more strong binders than Bernoulli on average. The improved performance is statistically significant (z=2.25, p=0.025), with a 95% confidence interval of [0.9 13.2].

**Table 3:** Mixed Linear Model - Quantum vs Bernoulli.

| Term | quantum sim vs Bernoulli | quantum real vs Bernoulli |
| --- | --- | --- |
| Baseline strong binders | 673.6 | 673.6 |
| Absolute improvement | 11.3 | 7.1 |
| Relative improvement | 1.7% | 1.0% |
| p-value | **<0.001** | **0.025** |

#### A.3.2 Comparison between Quantum and Gaussian

Table 4 reports the statistics of the difference in mean and standard deviation of the number of strong binders produced by the simulated and real quantum distributions compared to the Gaussian distribution. The results for both quantum distributions suggest a performance advantage in the mean number of strong binders, without achieving p < 0.05 significance. However, there is a significant difference between the variances of these distributions, with the real quantum distribution being significantly narrower according to both metrics and the simulated quantum distribution being significantly narrower according to the F-test.

**Table 4:**
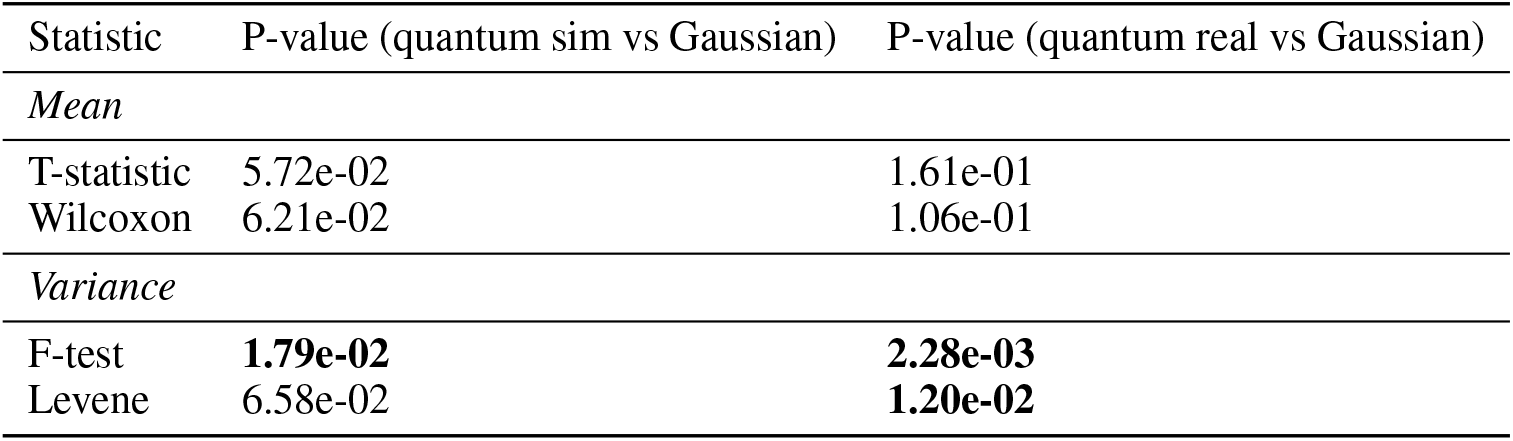
Statistics - Quantum vs Gaussian.

| Statistic | P-value (quantum sim vs Gaussian) | P-value (quantum real vs Gaussian) |
| --- | --- | --- |
| *Mean* |  |  |
| T-statistic | 5.72e-02 | 1.61e-01 |
| Wilcoxon | 6.21e-02 | 1.06e-01 |
| *Variance* |  |  |
| F-test | **1.79e-02** | **2.28e-03** |
| Levene | 6.58e-02 | **1.20e-02** |

Table 5 reports the results from the mixed linear model. On average, the Gaussian prior produces 674.3 strong binders out of 1000 generation attempts for each allele. The simulated quantum prior produces 10.6 more strong binders than the Gaussian prior on average. The performance improvement is statistically significant (z=3.31, p=0.001), with a 95% confidence interval of [4.3, 16.9]. The real quantum prior produces 6.3 more strong binders than Gaussian on average. The quantum effect is statistically significant (z=1.968, p=0.049), with a 95% confidence interval of [0.025, 12.667].

**Table 5:** Mixed Linear Model - Quantum vs Gaussian.

| Term | quantum sim vs Gaussian | quantum real vs Gaussian |
| --- | --- | --- |
| Baseline | 674.3 | 674.3 |
| Quantum improvement | 10.6 | 6.3 |
| Relative effect | 1.6% | 0.9% |
| p-value | **0.001** | **0.049** |

#### A.3.3 Discussion

Taken together, these results confirm that the two quantum priors lead to different overall results compared to the two baseline classical priors, with the clearest separation emerging in the allele-aware comparison. The higher mean number of strong binders achieved by the quantum priors, combined with the smaller variance of the results, shows that the quantum priors produce more consistent generative performance than the classical priors.

### A.4 Additional in-vitro results

In addition to testing 20 quantum-prior-generated peptides for each allele, we also tested 5 Gaussian-generated and 5 Bernoulli-generated peptides. Figure 9 shows in-vitro results including the best-performing classically-generated peptides. While the mean OD405 values were similar between the best-performing sets, the single strongest signal (maximum OD405) observed for each allele arose from a quantum-prior–generated peptide. This may be a consequence of the higher sequence diversity achieved by the quantum prior.

**Figure 9:**
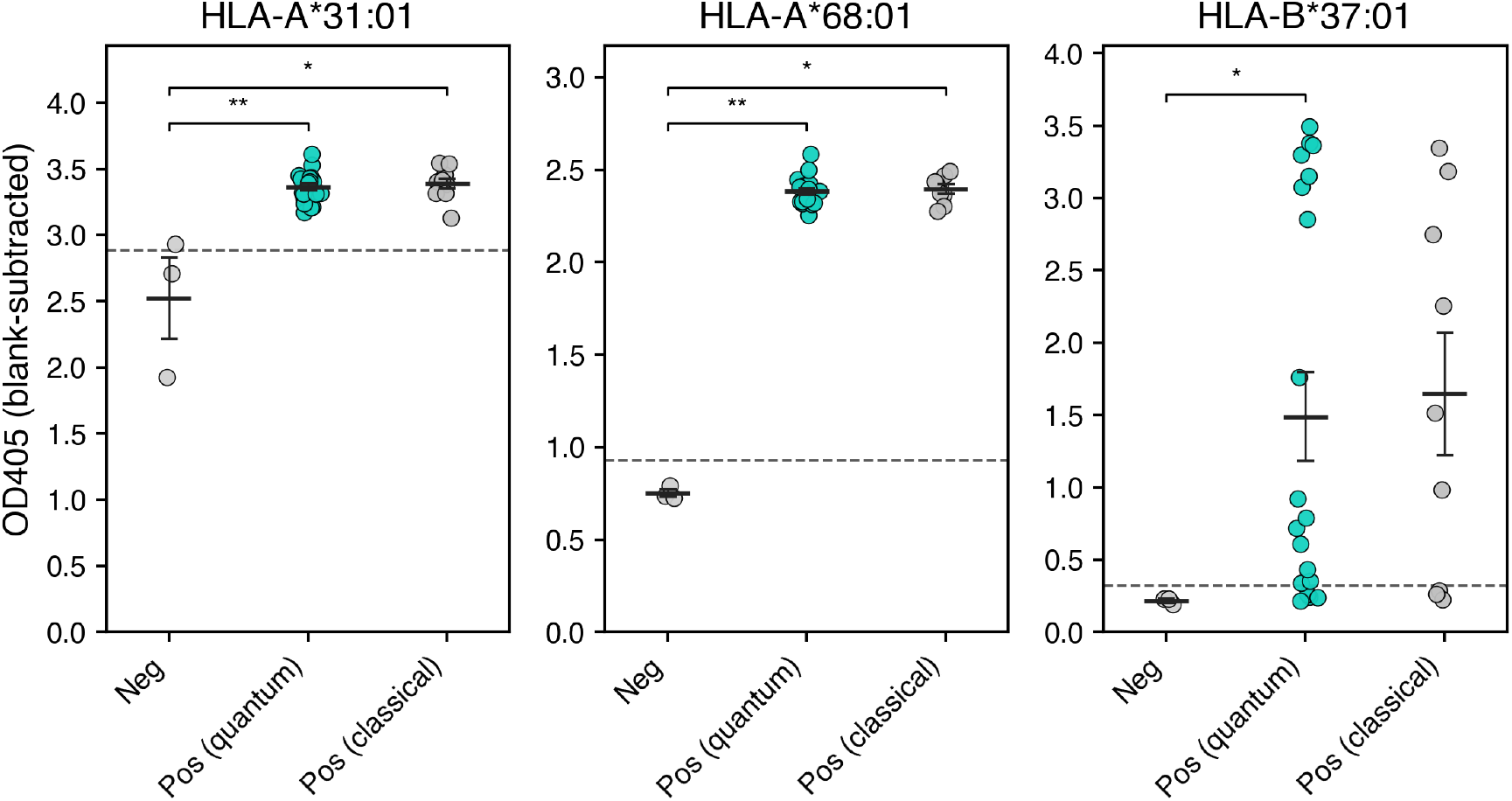
Peptide-dependent pMHC stability measured by sandwich ELISA. Points represent mean of technical triplicates; horizontal bars indicate mean ± s.d. The dashed line indicates the positivity threshold defined as mean of UV-only control without peptide.

### A.5 Cost of classically simulating photonic quantum processors

Though our work used photonic quantum processors with 32 channels, which is small enough to be exactly classically simulable, at larger scales beyond a few tens of photons and channels an exact classical simulation is no longer feasible. However, approximate methods for simulating a photonic quantum system exist. These approximate methods generally aim to reproduce the low-order statistics of a boson sampling distribution, making them practically hard to distinguish from the original quantum distribution. We expect these approximate methods would reach similar performance to the real quantum system in our experiments, in terms of the number of strong binders. However, their use incurs overheads in terms of wall clock time:

- The work in [43] involves sampling from a Boltzmann machine and has a runtime at least quadratic in the number of channels of the photonic system.
- The work in [44] requires 1 GPU per optical channel, and required about 10 minutes to initially set up the simulation of a given optical circuit
- The work in [45] involves initially calculating a large number of correlators before the sampling process can begin. Whereas a real hardware system they compare to generated 10 million samples in 10 seconds, their algorithm generated 10 million samples in 115 minutes.

To ensure an apples-to-apples comparison between the quantum distributions and the classical distributions, our current work does not attempt to train the quantum distribution, so even though these classical approximation methods may be slow it may still be feasible to pre-compute samples using them.

However, we argue that these approximate methods become impractical in more general settings when the parameters of the quantum processor are trained jointly with the neural network. Joint training of the prior distribution and the neural network often leads to improved results [15], and training methods exist for photonic quantum processors [56]. To enable joint training, samples from the prior distribution must be drawn at a similar rate to the rate at which they are consumed by the neural network. While photonic quantum processors can in principle operate at sufficiently high speeds, the classical approximation methods described above are likely to cause significant overheads in runtime that make them unusable in practice in this scenario.

### A.6 Detailed in vitro results

The HLA alleles used for in vitro validation (tables 6-8) were selected based on availability and low representation in the training dataset. MHC integrity was assessed using a UV-exchange ELISA, and peptide-dependent folding was quantified as blank-subtracted OD405. Signals were evaluated relative to a no-peptide UV control (mean + 3×SD). Rank of the predicted binding score was extracted from NetMHCpan-4.2[4].

**Table 6:**
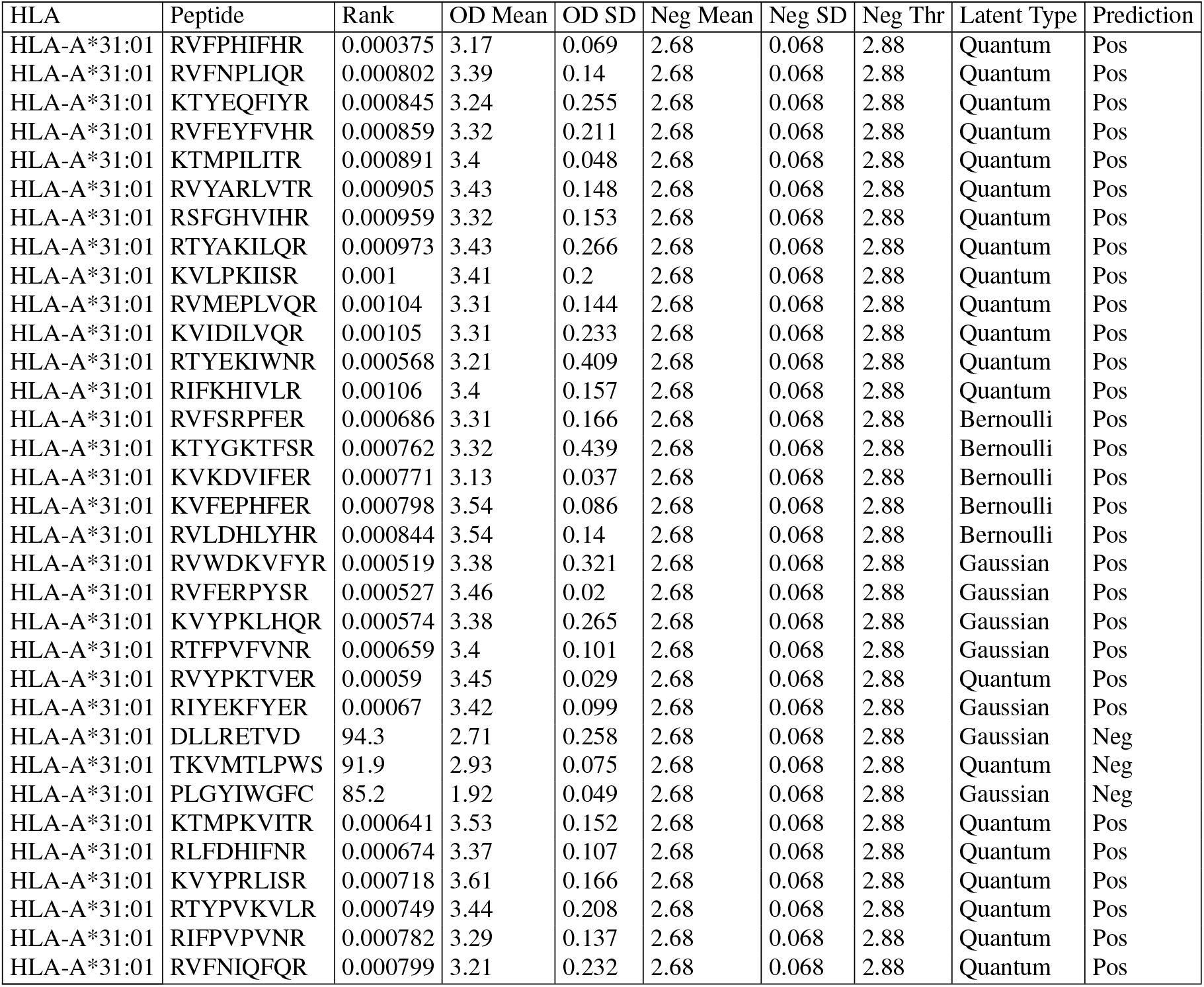
In vitro results for HLA-A*31:01.

**Table 7:**
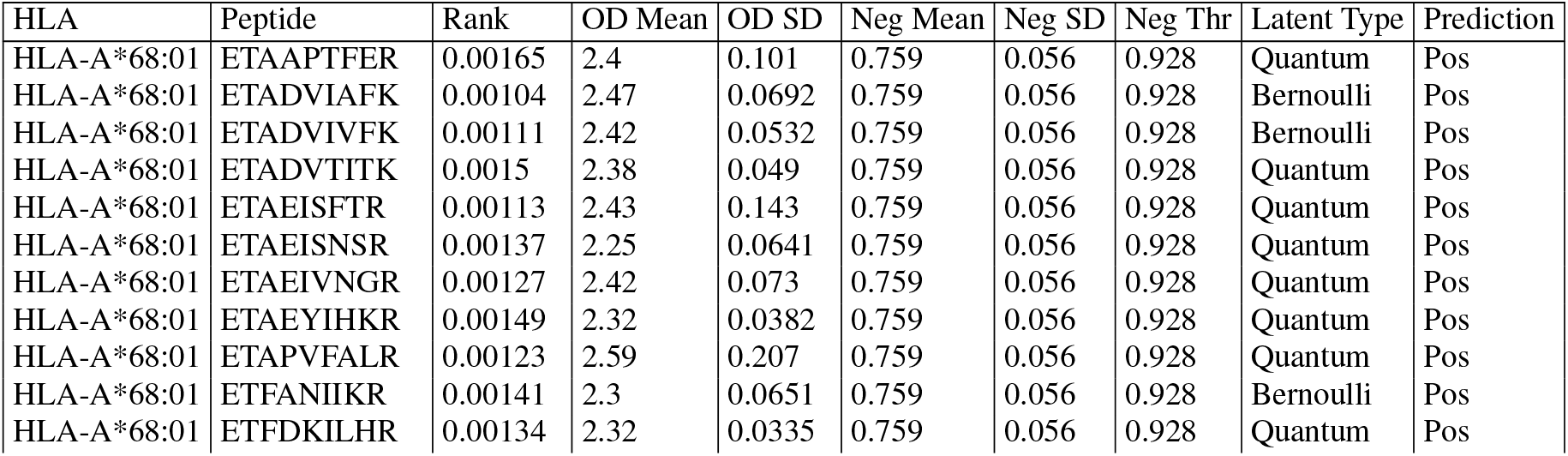
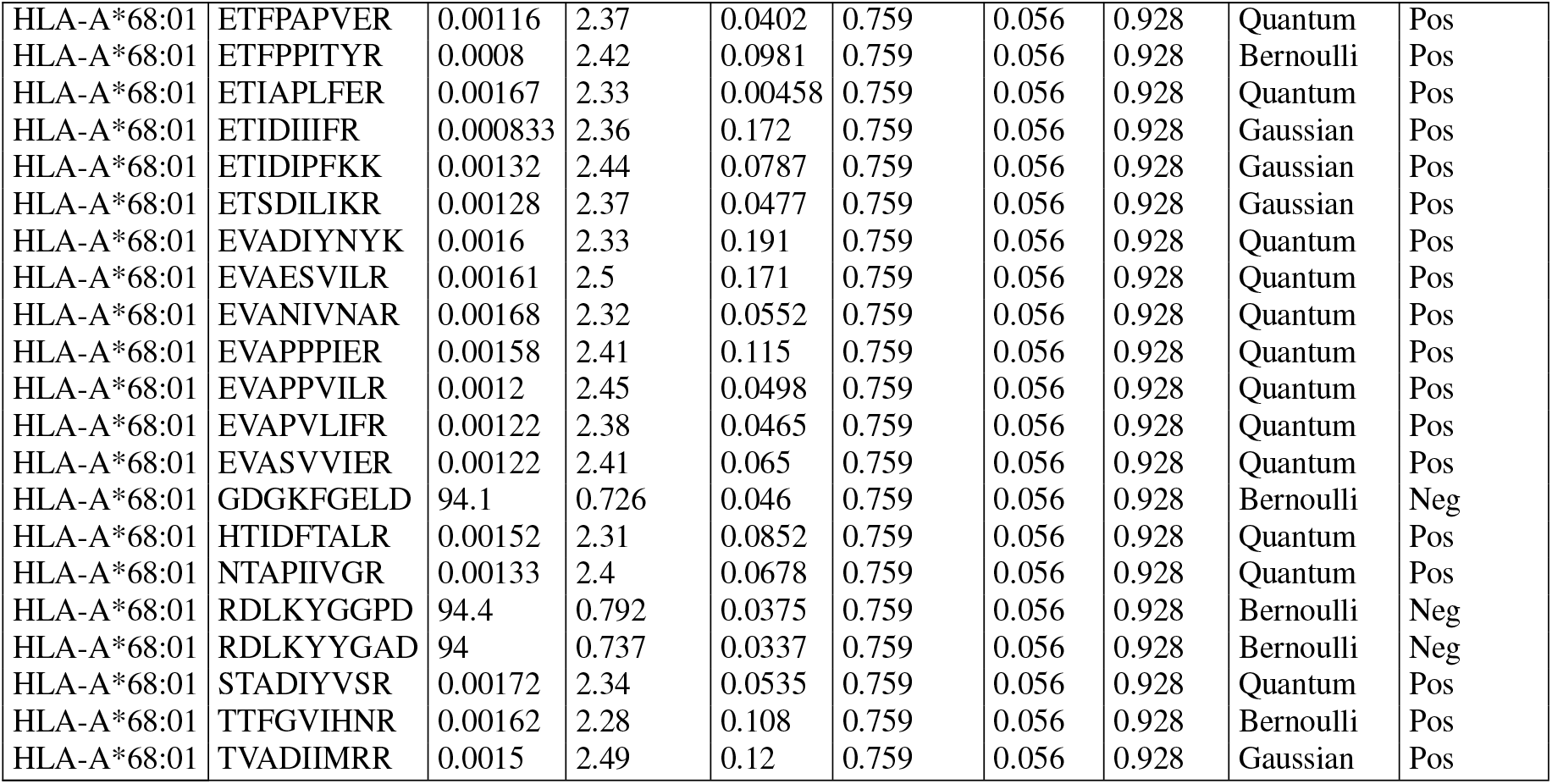
In vitro results for HLA-A*68:01.

**Table 8:**
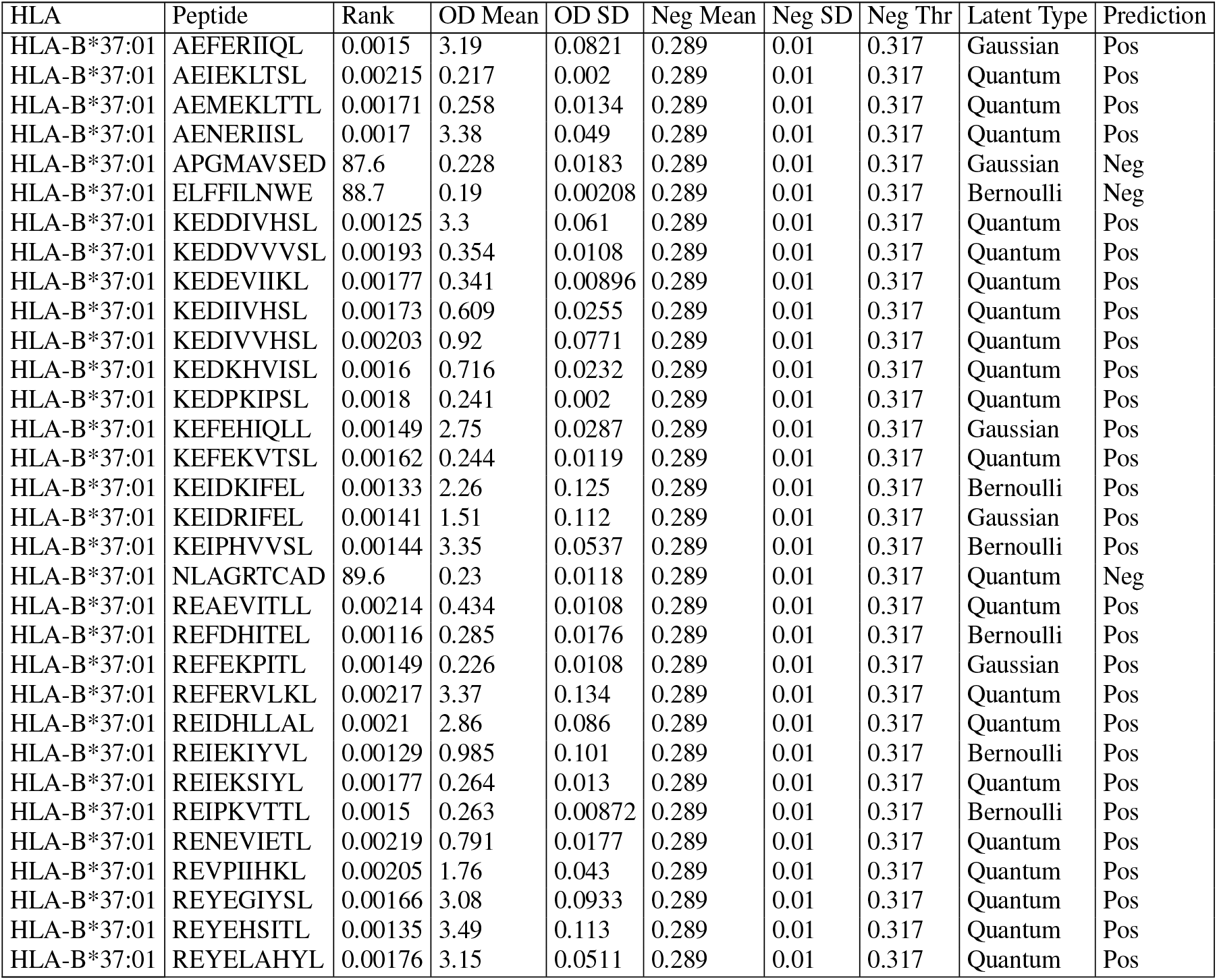
In vitro results for HLA-B*37:01.

